# Loss of CDC50A function drives Aβ/p3 production via increased β/α-secretase processing of APP

**DOI:** 10.1101/2020.11.12.379636

**Authors:** Marc D. Tambini, Luciano D’Adamio

## Abstract

The Amyloid Precursor Protein (APP) undergoes extensive proteolytic processing to produce several biologically active metabolites which affect Alzheimer’s disease (AD) pathogenesis. Sequential cleavage of APP by β- and γ-secretases results in Aβ, while cleavage by α- and γ-secretases produces the smaller p3 peptide. Here we report that in cells in which the P4-ATPase flippase subunit CDC50A has been knocked out, large increases in the products of β- and α-secretase cleavage of APP (sAPPβ/βCTF and sAPPα/αCTF, respectively) and the downstream metabolites Aβ and p3 are seen. These data indicate that APP cleavage by β/α-secretase are increased and suggest that phospholipid asymmetry plays an important role in APP metabolism and Aβ production.

## Introduction

Alzheimer’s disease (AD) is the most common form of dementia of the elderly[1]. Familial forms of AD are caused by mutations in the gene encoding the Amyloid Precursor Protein (APP) or in genes encoding Presenilin 1 or 2 (PS1/2), which process fragments of APP to produce plaque-forming Aβ peptides[2, 3]. APP is processed in two main pathways[4]. In the amyloidogenic pathway, APP undergoes sequential cleavage first by β-secretase (BACE1)[5] to produce one large soluble extracellular domain (sAPPβ) and one β-carboxy terminal fragment (βCTF). βCTF is then further processed by γ-secretase, composed in part by PS1 and PS2, to produce one molecule of Aβ and one molecule of APP-intracellular domain (AID/AICD)[6]. In the nonamyloidogenic pathway, APP is first cleaved within the Aβ region by α-secretase, to release a large soluble domain (sAPPα), and one αCTF. αCTF is then processed by γ-secretase to produce one molecule of the nonamyloidogenic p3 and one molecule of AID/AICD. Because metabolites of APP are biologically active and have numerous effects on AD-related histopathology and synaptic function[7, 8], the cellular processes and genes that govern APP processing are of considerable scientific interest.

When considering non-Mendelian, late-onset forms of AD, one of the themes to emerge from the large scale genome-wide association studies is that genes involved in lipid homeostasis[9] are responsible for, as non-deterministic genetic risk factors, the more common forms of AD. The classical and most well-validated example is Apolipoprotein E (ApoE)[10], an extracellular lipid transporter whose AD-associated isotype can significantly increase AD risk. The phosphatidylserine (PtdSer) sensing ability of the microglial protein Triggering Reception in Microglia 2 (TREM2) has been shown to be altered by AD-causing *TREM2* mutations[11]. The PtdSer transporter ATP-binding cassette A7 (ABCA7) likewise is mutated in late-onset AD[12].

Recently, cell division cycle protein 50A (CDC50A, also termed TMEM30A), a subunit of a class of lipid transporters called flippases[13], which transport phospholipids from the exofacial leaflet of the plasma membrane to the cytosolic face, has been reported to bind βCTF and affect APP-CTF levels in an *in vitro* overexpression model[14]. Given the constitutively active nature of PtdSer flippase activity, it is possible that an over expression paradigm may not reveal the true impact of CDC50A on APP processing, in addition to the possible confounding effects caused by overexpression artifacts. Indeed, the loss of CDC50A function, and therefore a loss of flippase function, has been shown to have a neurodegenerative phenotype in conditional KO models[15, 16], we wished to determine the effect of loss of CDC50A function has on APP processing. Here, we characterize APP processing in a human CDC50A knockout (KO) tumor cell line. We report a significant increase in p3 and Aβ production in CDC50A-KO cells, driven by increase in α- and β-processing of APP and other substrates.

## Results

### CDC50A loss of function increases Aβ, p3, and sAPP secretion in cultured cells

CDC50A-KO cells were generated by Horizon Discovery via the CRISPR/Cas9-mediated 16-bair pair deletion in the *CDC50A* coding region of parental wild-type HAP1 cells. RT-PCR of CDC50A RNA demonstrates an 18-fold decrease in CDC50A transcript in CDC50A-KO cells, likely the result of nonsense-mediated decay (Fig 1A). Functional knockout of CDC50A was confirmed by testing surface Annexin V binding, an indirect measure of surface levels of PtdSer. HAP1 cells showed no Annexin V staining, consistent with intact PtdSer flippase activity (Fig. 1B, top panel), whereas CDC50A-KO cells were positive (Fig. 1B, middle panel), indicating increased surface levels of PtdSer. HAP1 cells could be made to robustly expose PtdSer by induction of apoptosis with staurosporine treatment (Fig. 1C, bottom panel). These data indicate that CDC50A-KO cells reproduce the expected loss of function of PtdSer flippase activity.

**Figure 1.**
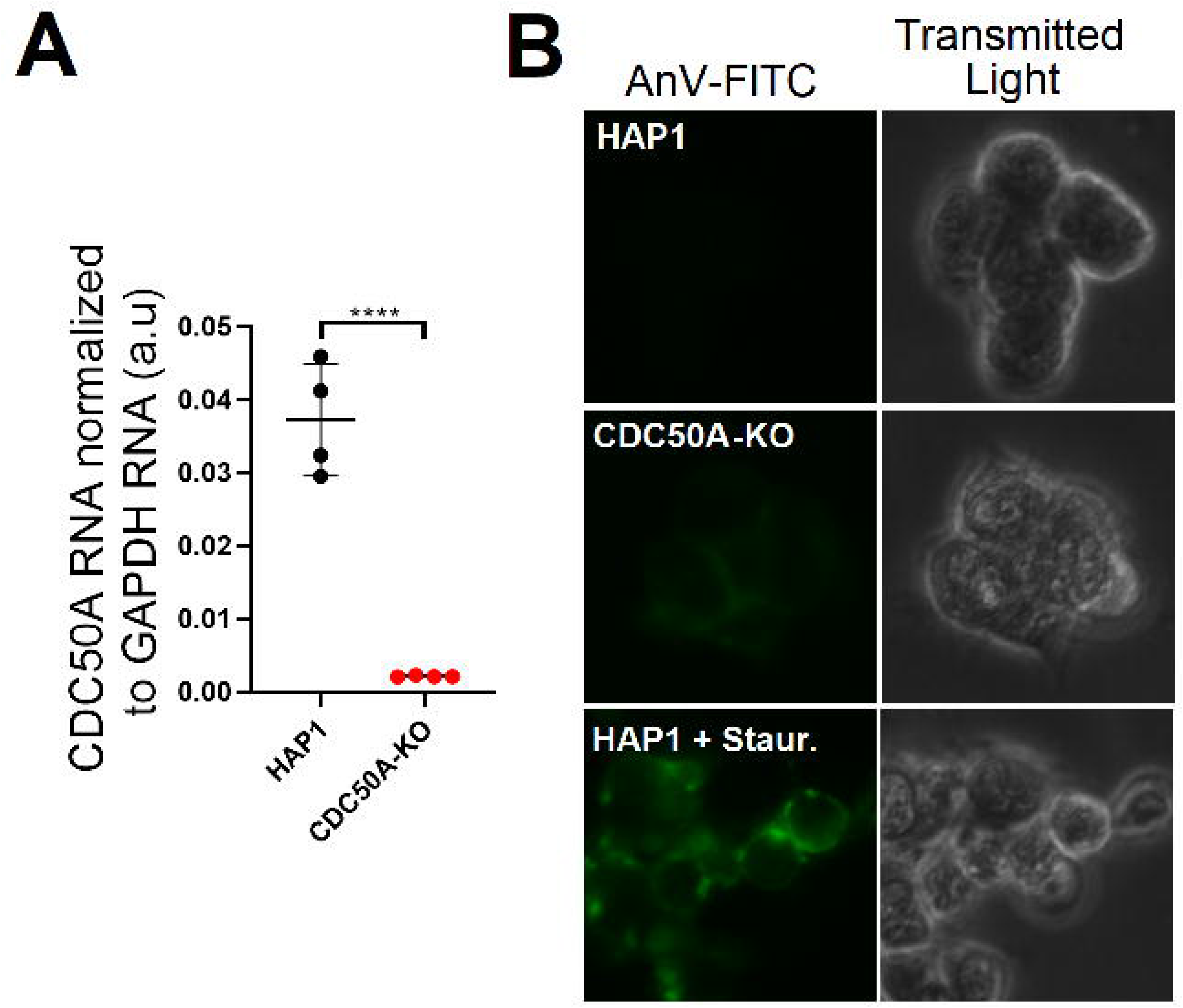
Validation of CDC50A-KO by RT-PCR and PtdSer binding assay. A.) CDC50A RNA was detected in wild-type HAP1 cells and CDC50A-KO cells. Data were analyzed by two-sided Student’s *t*-test and represented as mean ± SD. n=4 per genotype, ****p<0.0001. B.) Annexin V-FITC (AnV-FITC) binding was visualized in HAP1, CDC50A-KO, and staurosporine-treated (1μM, 2hr) HAP1 cells by fluorescent microscopy.

APP metabolite levels of HAP1 and CDC50A-KO cell conditioned media were determined by ELISA. Two different ELISAs were used to determine Aβ levels: one with the detection antibody 4G8, which recognizes Aβ amino acids 3-8, and one with detection antibody 6E10, which recognizes Aβ amino acids 17-24. 4G8 therefore detects Aβ and p3, the fragment of APP-CTF that is produced from α- and γ-cleavage, whereas 6E10 only detects Aβ. The difference in 4G8 and 6E10 signal can then be used to calculate the level of p3. Note that in brain homogenates, it has been found that the differing activities of 6E10 and 4G8 are such that p3 cannot be reliably estimated by this method[17], whereas in cultured cells, this method has been shown to be reliable[18, 19]. In cultured media from CDC50A-KO cells, a 9-fold increase in Aβ40 is seen, as detected by 6E10, and a 26-fold increase in the calculated p3(Aβ17-40) as compared to HAP1 conditioned media (Fig. 2A). The longer Aβ42 and p3(Aβ17-42) are also increased in CDC50A-KO conditioned media, at 10-fold and 3-fold respectively (Fig. 2B). Levels of the large soluble ectodomains of APP derived from the α- and β-cleavage of APP, termed sAPPα and sAPPβ respectively, were determined by ELISA of HAP1 and CDC50A-KO conditioned media. Both sAPPα and sAPPβ were found to be significantly increased, at 8-fold and 3-fold, in media from CDC50A-KO cells as compared to controls. Overall, these data indicate the increased α- and β-processing of APP in CDC50A-KO cells.

**Figure 2.**
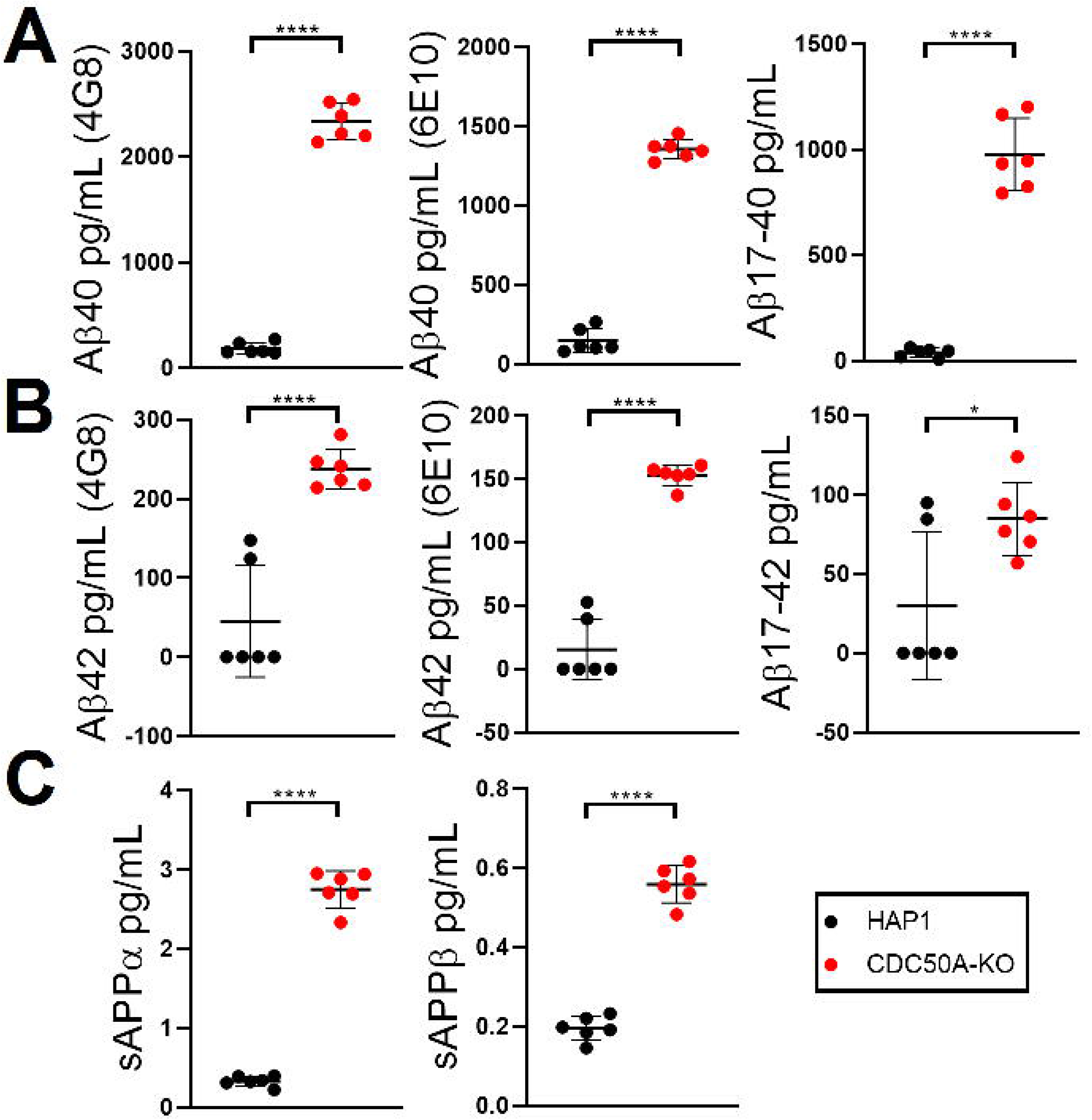
ELISA of APP metabolites in WT and CDC50A-KO cell conditioned media. A.) Levels of Aβ40 were determined by ELISAs with either 4G8 (left graph, detects both p3 and Aβ) or 6E10 (middle graph, only detects Aβ) detection antibodies. p3/Aβ17-40 (right graph) was calculated by subtracting 4G8 values from 6E10 values. B.) Levels of Aβ42 were determined by ELISAs with either 4G8 (left graph, detects both p3 and Aβ) or 6E10 (middle graph, only detects Aβ) detection antibodies. p3/Aβ17-42 (right graph) was calculated by subtracting 4G8 values from 6E10 values. C.) Levels of sAPPα and sAPPβ were determined by ELISA. For all ELISAs, data were analyzed by two-sided Student’s *t*-test and represented as mean ± SD. n=6 per genotype, *p<0.05, ****p<0.0001.

### CDC50A-KO cells show general altered levels of γ-secretase substrates under γ-secretase inhibition

Given that increased Aβ and p3 levels can be the result of increased γ-secretase activity or increased total substrate levels of the full length APP, we wished to determine membrane bound APP and APP-CTF levels of HAP1 and CDC50A-KO lysates by Western analysis. While a 54% and 57% increase in immature and mature APP were noted in CDC50A-KO cells (Fig. 3A), this increase is not of a sufficient magnitude to explain the manifold increase in Aβ and p3 levels secreted by CDC50A-KO cells. APP-CTFs are likewise unaltered by CDC50A knockout (Fig. 3A). We tested another substrate of γ-secretase, N-cadherin, and found no difference in the levels of full-length N-cadherin and a minor increase in N-cadherin-CTFs (Fig. 3B).

**Figure 3.**
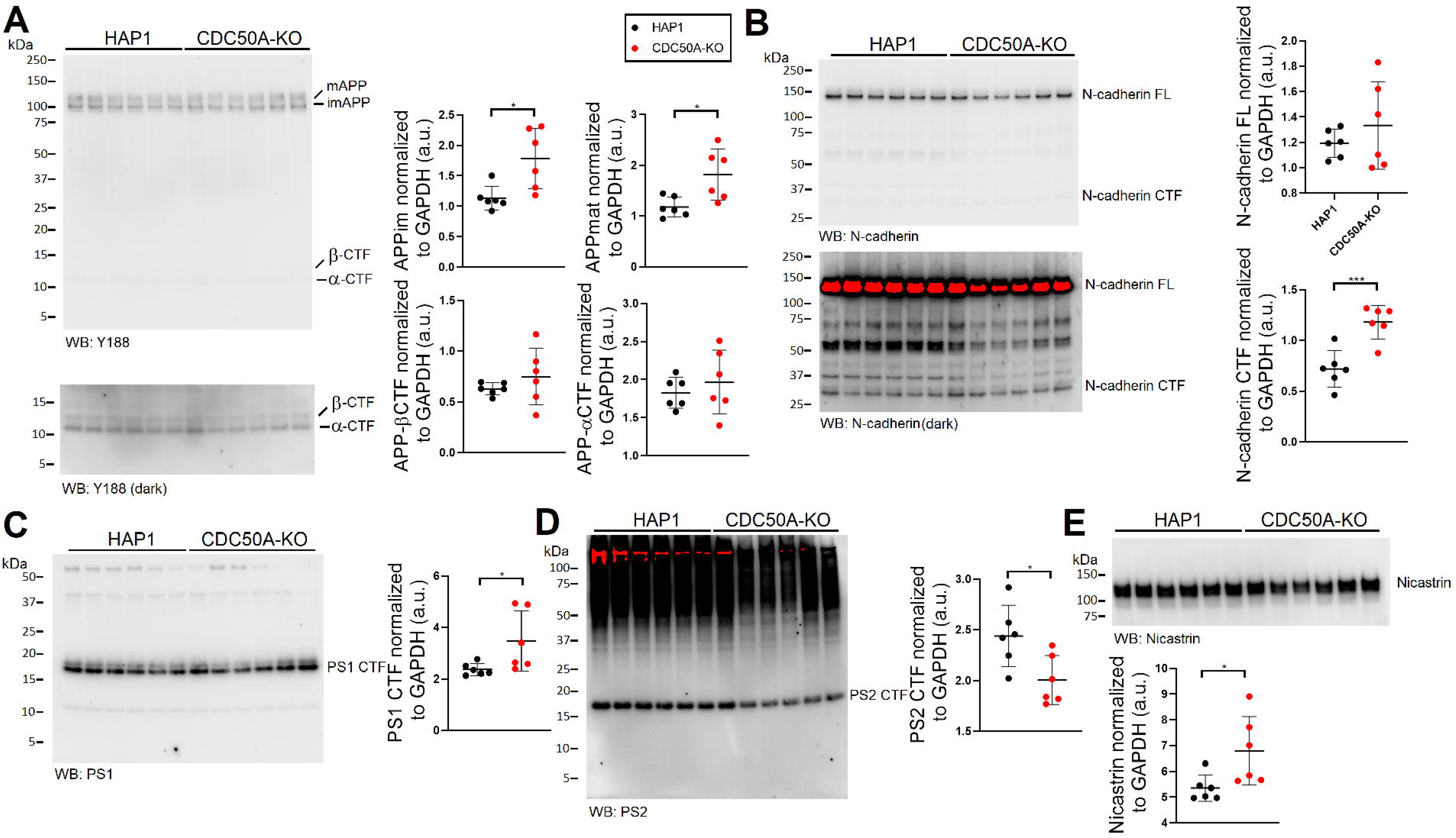
Western analysis of APP, N-cadherin, and γ-secretase component levels of WT and CDC50A-KO cell lysates. The following antibodies were used to probe wild-type HAP1 and CDC50A-KO cell lysates: A.) Y188 APP C-terminus. Mature and immature full-length APP, and APP-CTFs, both αCTF and βCTF, are indicated. APP-CTFs were quantitated from the darker exposure. B.) N-cadherin C-terminus. Full length N-cadherin and N-cadherin-CTFs are indicated. Full-length N-cadherin was quantitated from the lighter exposure, and N-cadherin-CTFs were quantitated from the darker exposure. Note the band designated as N-cadherin CTF has been experimentally tested (Fig. 4B) and identified as N-cadherin CTF via its sensitivity to γ-secretase inhibition. All other bands are non-specific signals. C.) PS1 C-terminus. PS1-CTF is indicated, with no full length PS1 detected. D.) PS2 C-terminus. PS2-CTF is indicated, with no full length PS2 detected. E.) Nicastrin. Quantitation is shown at the right or below each blot. All data were normalized to GAPDH signal, analyzed by two-sided Student’s t-test, and represented as mean ± SD. n=6 per genotype, *p<0.05, ***p<0.001.

Components of γ-secretase itself showed minor alterations, with CDC50A-KO cells having 32% more PS1-CTF levels (Fig. 3C), 22% lower PS2-CTF levels (Fig. 3D), and 27% increased levels of nicastrin (Fig. 3E). There is no evidence of altered PS1 or PS2 auto-catalysis (Fig. 3C-D), as no full-length PS1 or PS2 is detectable. The data do not support the idea that the changes in Aβ and p3 is the result of altered γ-secretase function.

sAPPα and sAPPβ are better indicators of α- and β-cleavage of APP than levels of αCTF and βCTF, which undergo further processing by various pathways. We wished to determine if the increase in αCTF and βCTF that are predicted by the increases in sAPPα and sAPPβ can be revealed by blocking APP-CTF degradation with a γ-secretase inhibitor. HAP1 and CDC50A-KO cells were treated with compound E, a potent, selective, noncompetitive inhibitor of γ-secretase. Compound E activity was confirmed by the large increase in APP-CTFs seen by Western analysis (Fig. 4A). Interestingly, under γ-inhibition, CDC50A-KO cell lysates had 15-fold higher αCTF and 44-fold higher βCTF than wild-type HAP1 cells (Fig. 4A). This effect is not specific to APP, as N-cadherin-CTFs were also significantly increased in γ-inhibited CDC50A-KO cells under γ-inhibited controls (Fig. 4B). The absence of any difference in APP-CTFs or N-cadherin-CTFs in CDC50A-KO cells in the steady state indicates that γ-secretase can efficiently processes the increased substrates that result from elevated α- and β-activity.

**Figure 4.**
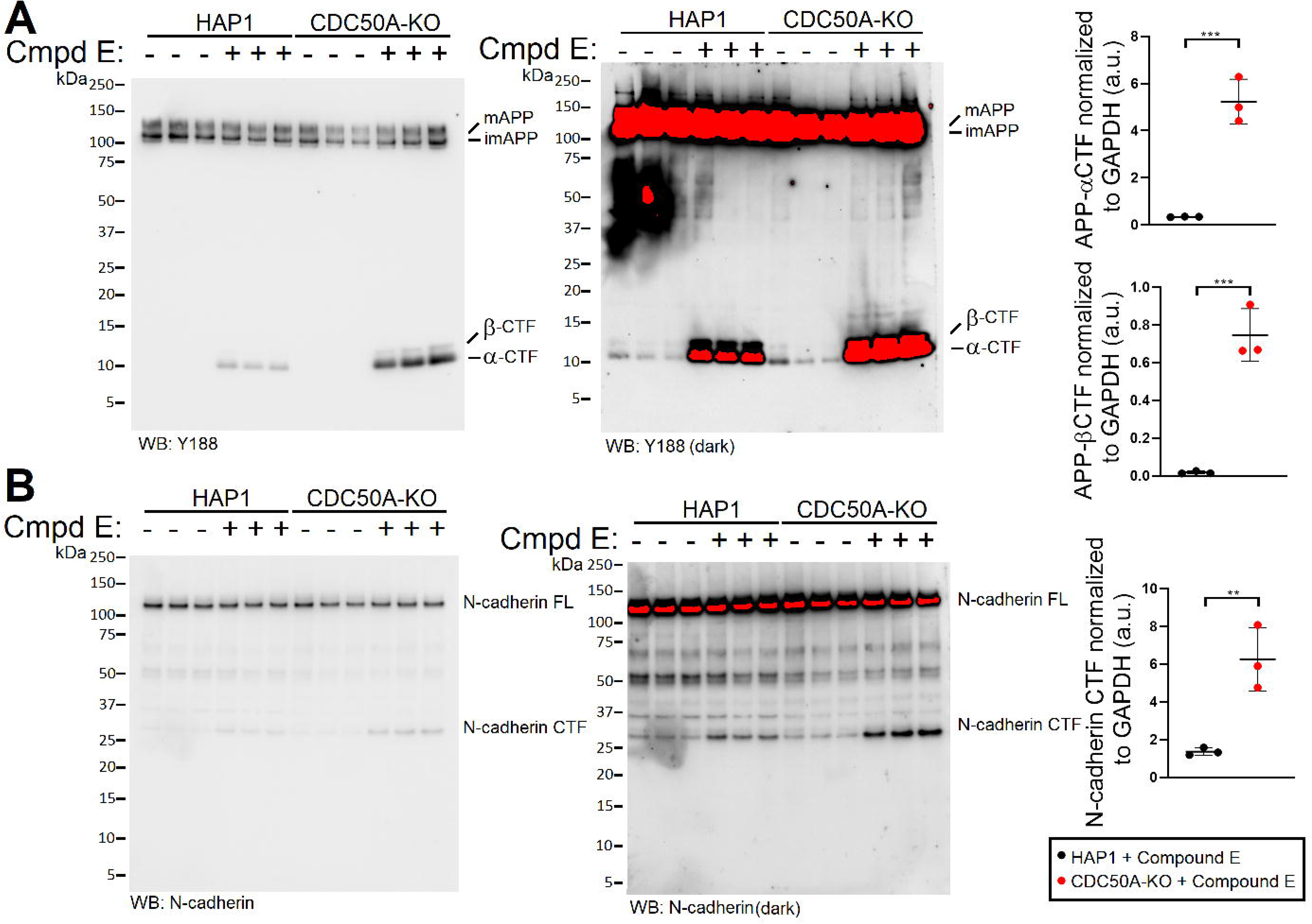
Western analysis of APP and N-cadherin in WT and CDC50A-KO cell lysates under γ-secretase inhibition. Wild-type HAP1 and CDC50A-KO cells were either untreated or treated with compound E, a potent, selective, non-competitive γ-secretase inhibitor. Cell lysates were probed as follows: A.) Y188 APP C-terminus. Mature and immature full-length APP, and APP-CTFs, both αCTF and βCTF, are indicated. APP-CTFs were quantitated from the γ-secretase inhibited samples using the lighter exposure. A darker exposure indicates the degree of γ-secretase inhibition of compound E treated samples. B.) N-cadherin C-terminus. Full length N-cadherin and N-cadherin-CTFs are indicated. N-cadherin-CTFs from the γ-secretase inhibited samples were quantitated from the darker exposure. All data were normalized to GAPDH signal, analyzed by two-sided Student’s *t*-test, and represented as mean ± SD. n=3 per genotype, **p<0.01, ***p<0.001.

## Discussion

In this study, we demonstrate that APP undergoes increased α- and β-cleavage to produce more p3 and Aβ in CDC50A-KO cells. The mechanism of this increased processing is unclear. Loss of membrane phospholipid asymmetry has been reported to have numerous effects on endocytic trafficking. We speculate that this disrupted trafficking would result in the general altered processing of membrane proteins. APP, a type 1 transmembrane protein, is generated in the ER, travels along the biosynthetic pathway to the plasma membrane, and is then endocytosed to late endosome/lysosome. Along this pathway, it is subject to numerous cleavages. At the plasma membrane, APP is predominantly cut by α-secretase, while the acidic lumen of the late endosome/lysosome favors β-cleavage. Given that both the amyloidogenic and non-amyloidogenic pathways are upregulated in CDC50A-KO cells, it seems unlikely phospholipid asymmetry causes APP mislocalization, or selective organellar dysfunction. Instead, it is possible that there is a generalized increase in membrane turnover, and this idea is supported by the observation that another type-1 transmembrane protein, N-cadherin, was affected in a similar manner.

## Materials and Methods

### Cell Lines and reagents

Wild-type HAP1 and CDC50A-KO cell lines were purchased from Horizon Discovery (C631, HZGHC005423c009). Cells were cultured in DMEM (Gibco 11995-065), supplemented with 10% FBS (Gibco 26140-079) and 1% antibiotic–antimycotic (Gibco 15240) at 37°C and 5% CO_2_. Compound E (EMD Millipore 565790) was used at a final concentration of 10μM for 18h for the γ-secretase inhibited samples. Staurosporine (Cell Signaling 9953) was used at final concentration of 1μM for 2hr to induce apoptosis.

### Western Analysis

Cells were grown in a monolayer to 90% confluency, washed 3x in PBS, and scraped off into ice-cold homogenization buffer (20 mM Tris-base pH 7. 4, 250 mM sucrose, 1 mM EDTA, 1 mM EGTA) plus protease/phosphatase inhibitors (ThermoScientific). Cells were lysed via sonication and total protein was measured by Bradford analysis. 10 μg of protein plus LDS Sample buffer/10% β-mercaptoethanol (Invitrogen NP0007) were separate by PAGE on a 4–12% Bis-Tris polyacrylamide gel (Biorad 3450125), transferred onto nitrocellulose using the Trans-blot Turbo system (Biorad) and visualized by red Ponceau staining. After membranes were blocked in 5%-milk (Biorad 1706404), the following primary antibodies were applied overnight at 4°C, at 1:1000 dilution in blocking solution (Thermo 37573): APP-Y188 (Abcam ab32136), PS2 (Cell signaling, 2192), Nicastrin (Cell signaling, 5665), N-cadherin (Cell signaling, 14215), PS1 (Cell signaling, 5643) and GAPDH (Sigma G9545). Either anti-mouse (Southern Biotech 1031-05) or a 1:1 mix of anti-rabbit (Southern Biotech, OB405005) and anti-rabbit (Cell Signaling, 7074), were diluted 1:1000 in 5% milk and used against mouse and rabbit primary antibodies for 30 min, RT, with shaking. Blots were developed with West Dura ECL reagent (Thermo, PI34076) and visualized on a ChemiDoc MP Imaging System (Biorad). Signal intensity was quantified with Image Lab software (Biorad). Data were analyzed using Prism software and represented as mean ± SD GAPDH (Sigma G9545). After extensive washings in PBS/Tween20 0. 05%, the following secondary antibodies were used diluted 1:1000 in 5%-milk: anti-mouse (Southern Biotech, 1031–05) and a 1:1 mix of anti-rabbit (Southern Biotech, 4050– 05) and anti-rabbit (Cell Signaling, 7074). Secondary antibodies were incubated with membranes for 30 min, RT, with shaking). After extensive washings in PBS/Tween20 0. 05%, blots were developed with West Dura ECL reagent (Thermo, PI34076), visualized with ChemiDoc MP Imaging System (Biorad) and signal intensities were quantified with Image Lab software (Biorad).

### ELISA

Measurement of Aβ40, Aβ42, sAPPα, and sAPPβ content of conditioned media was preformed using the following Meso Scale Discovery: PLEX Plus Aβ Peptide Panel 1 6E10 (K15200G), V-PLEX Plus Aβ Peptide Panel 1 4G8 (K15199G), sAPPα/sAPPβ (K15120E), according to the manufacturer’s recommendations. Fresh media was applied to cells for 24 hours, and dead cells were removed by a 10m, 20,000 x g spin before the supernatant was used for analysis. Plates were read on a MESO QuickPlex SQ 120. Data were analyzed using Prism software and represented as mean ± SD.

### RT-PCR

RNA was extracted from cultured cells using the RNeasy RNA Isolation kit (Qiagen 74104) and used as template for cDNA generation with a High-Capacity cDNA Reverse Transcription Kit (Thermo 4368814). Real-time PCR was performed using 50 ng of cDNA and TaqMan Fast Advanced Master Mix (Thermo 4444556). GAPDH mRNA was detected with probe Hs02786624_g1 and CDC50A mRNA was detected with probe Hs01092148_m1 (Thermo). Samples were analyzed on a QuantStudio 6 Flex Real-Time PCR System (Thermo). LinRegPCR software (hartfaalcentrum.nl) was used to quantify relative mRNA amounts.

### Annexin V labeling and microscopy

Cells were grown on glass coverslips, washed 3x in PBS and stained with Annexin V-FITC on ice, according to the manufacturer’s recommendations (BD Pharmingen 550911). Cells were fixed with 4% paraformaldehyde for 15m at room temperature, washed, mounted onto glass slides and visualized using an Axiovert fluorescence microscope (Zeiss).

### Statistical Analysis

Statistical significance was evaluated using the two-sided Student’s *t*-test. Statistical analysis was performed with GraphPad Prism v8 for PC. Significant differences were accepted at p < 0.05.

---

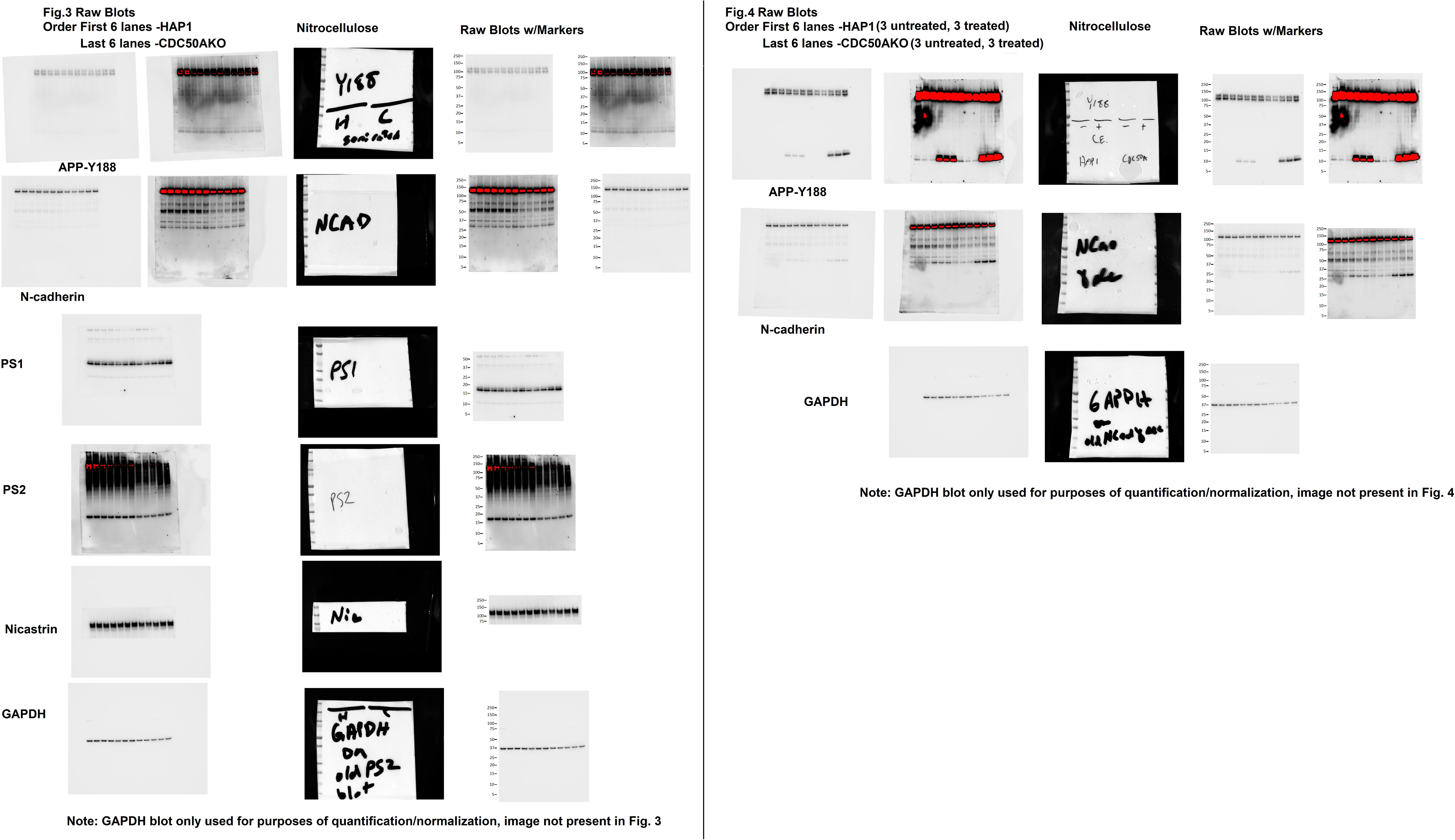

